# Uncovering the Pathophysiological Pattern of Expression from Integrated Analysis across Uniformly Processed RNA Sequencing COVID-19 Datasets

**DOI:** 10.1101/2025.06.18.660454

**Authors:** Andrew Cole, Bryce Fukuda, Thomas Dahlstrom, Ling-Hong Hung, Ka Yee Yeung

## Abstract

**Background:** Post-acute sequelae of SARS-CoV-2 infection (PASC) affects millions globally, yet the molecular mechanisms underlying acute COVID-19 and its chronic sequelae remain poorly understood.

**Methods:** We performed an integrative transcriptomic analysis of three independent RNA-seq datasets, capturing the complete COVID-19 pathophysiology from health through acute severe infection to post-acute sequelae and mortality (n=142 total samples). We implemented a containerized analytical pipeline from data download, quantification, differential gene expression to uniformly process these three RNA-seq datasets.

**Results:** Our analysis reveals striking molecular dichotomies contrasting disease phases with profound clinical implications. Acute severe/critical COVID-19 reveals predominant enrichment of TNF-α signaling via NF-κB pathways (normalized enrichment score >2.5, FDR <0.001), reflecting a cytokine storm pathophysiology characterized by rapid inflammatory developments involving IL-6, TNF-α, and anti-apoptotic responses. In contrast, PASC patients exhibit dominant enrichment of Myc Targets V1 and Oxidative Phosphorylation pathways (NES >2.2, FDR <0.005), indicating important shifts toward cellular adaptation. Pathway signature analysis identifies core differentially expressed genes that reliably distinguish disease phases, thereby offering objective biomarkers for precision diagnosis and monitoring.

**Conclusions:** These findings establish a comprehensive molecular framework distinguishing acute inflammatory from chronic metabolic COVID-19 phases, with potential clinical applicability. TNF-α/NF-κB pathway signatures identify patients at risk for severe disease progression, while Myc/OXPHOS signatures allow objective PASC diagnosis, addressing current reliance on subjective and eliminative diagnosis. This integrative analytical framework has utility beyond COVID-19, offering an applicable approach for precision medicine implementation across other diseases processes.

**Clinical Significance:** This study transforms COVID-19 from a symptom-based to a molecularly-defined disease spectrum, enabling precision diagnosis, prognostic monitoring, classification, and targeted therapeutic possibilities based on pathway-specific biomarkers rather than subjective clinical assessments.

## Introduction

Post-acute sequelae of SARS-CoV-2 infection (PASC), also known as Long COVID, has emerged as one of the most pressing healthcare challenges, affecting an estimated 6 in 100 individuals following acute COVID-19 infection^1^. It encompasses a diverse constellation of symptoms that include fatigue, cognitive impairment, dyspnea, and multi-organ dysfunction that can persist for months or years after initial infection with SARS-CoV-22.Despite extensive clinical characterization along with the development of diagnostic criteria, the molecular mechanisms underlying PASC remain poorly understood, blurring objective diagnostic biomarkers^3^.

The heterogeneous nature of PASC presents unique challenges with patients exhibiting varying symptom severity, duration, and organ system involvement, suggesting distinct underlying pathophysiological mechanisms. It is also notable that the relationship between acute COVID-19 severity and subsequent PASC development remains complex and incompletely characterized^4^. While some patients with COVID-19 infection subsequently develop severe long-term sequelae, others recover completely without evidence of persistent PASC related signs and symptoms^5^. This clinical heterogeneity necessitates comprehensive molecular investigation that can capture the full spectrum of disease manifestations to identify common versus distinct mechanistic pathways related to active COVID-19 and PASC.

High-throughput transcriptomic profiling is now established as a powerful tool for dissecting complex disease mechanisms, with RNA sequencing (RNA-seq) providing transcriptome-wide insights into cellular responses and regulatory networks. The plethora of COVID-19 and PASC transcriptomic datasets in multiple repositories present attractive opportunities for understanding the genetic basis of disease pathophysiology^6^. However, realizing this potential involves addressing significant analytical challenges such as: heterogeneous experimental protocols, diverse computational pipelines, and technical batch effects that can obscure true biological signals^7^.

A critical limitation in current PASC research is the scarcity of studies that comprehensively capture the complete pathophysiological trajectory from health through acute infection to post-acute sequelae and, in severe cases, mortality. Most investigations focus on isolated parts of disease phases or compare limited patient subgroups, preventing identification of molecular signatures that distinguish disease progression pathways^8,9^. Additionally, the use of disparate analytical methodologies across studies introduces workflow variability that may complicate cross-study comparisons and meta-analyses.

Several groups have conducted integrative analyses of COVID-19 related transcriptomic data, each revealing important but incomplete insights into disease mechanisms. Gardinassi et al. performed a meta-analysis of multiple COVID-19 datasets focusing on immune cell populations, thereby identifying common interferon-stimulated gene signatures across acute infection studies^10^. Similarly, Arunachalam et al. integrated cross-sectional transcriptomic and proteomic data to characterize immune responses during acute COVID-19, revealing distinct trajectories associated with disease severity^11^. However, these studies primarily focused on acute infection phases and did not examine PASC which is an extension the post-resolution signs and symptoms from the initial COVID-19 infection.

More recently, researchers have investigated PASC-specific molecular signatures. Phetsouphanh et al., for example, identified persistent immune activation and complement dysregulation in patients with PASC, while Schultheiß et al. reported mitochondrial dysfunction and altered cellular metabolism^8,9^. Despite these advances, no study has systematically integrated and compared molecular signatures across the complete COVID-19 disease spectrum using uniformly processed datasets.

The challenge of integrating heterogeneous omics datasets has also been recognized across multiple disease contexts. Ritchie et al. developed frameworks for cross-platform genomics integration, while Love et al. established best practices for RNA-seq differential expression analysis^12,13^. These advances have availed the foundation for robust integrative analyses but require careful implementation to address study-specific variations.

To address these limitations, we developed an integrative analytical approach to uniformly process and analyze transcriptomic data across the complete COVID-19 pathophysiological spectrum. Our study design integrates three independent RNA-seq datasets: (1) Ryan et al., examining healthy controls and PASC patients with varying symptom severity from Australia^14^; (2) Yin et al., comparing individuals who recovered from COVID-19 with and without long-term sequelae from the United States^15^; and (3) Vlasov et al., profiling survivors and fatal cases of acute severe COVID-19 from Russia^16^. This comprehensive compendium captures the molecular landscape from health through acute severe infection to chronic post-acute sequelae and mortality.

We implemented a standardized, containerized, analytical pipeline that uniformly processes raw sequencing data through alignment, quantification, and differential expression analysis which eliminated technical confounders that could obscure biological insights. This approach enables direct molecular comparisons across disease states, datasets, and geographical populations while maintaining full analytical reproducibility.

## Materials and Methods

### Dataset Selection and Study Design

The datasets included in this integrative compendium were strategically selected to capture the complete pathophysiological eventualities of COVID-19, from health through acute infection to post-acute sequelae and mortality. This integrative approach was necessitated by the lack of studies containing samples representing all disease phases in a single study. We conducted extensive literature searches aligning to our study design using PubMed, Google Scholar, and AI aided semantic/transcriptomic searches to identify suitable RNA-seq datasets that collectively would enable investigation of our three primary research objectives: (1) characterizing gene expression pathways of dominance in acute severe COVID-19, (2) identifying and characterizing the gene expression signatures in PASC, and (3) identifying and profiling molecular biomarkers for phase-specific disease classification.

Our final compendium integrated three independent studies from different continents, ensuring geographic diversity and robust representation across disease states: Ryan et al. (Australia) containing healthy controls and PASC patients with varying symptom severity; Yin et al. (United States) comparing recovered individuals from COVID-19 with and without long-term sequelae; and Vlasov et al. (Russia) profiling survivors versus fatal cases from acute severe COVID-19, Figure 1.

**Figure 1.**
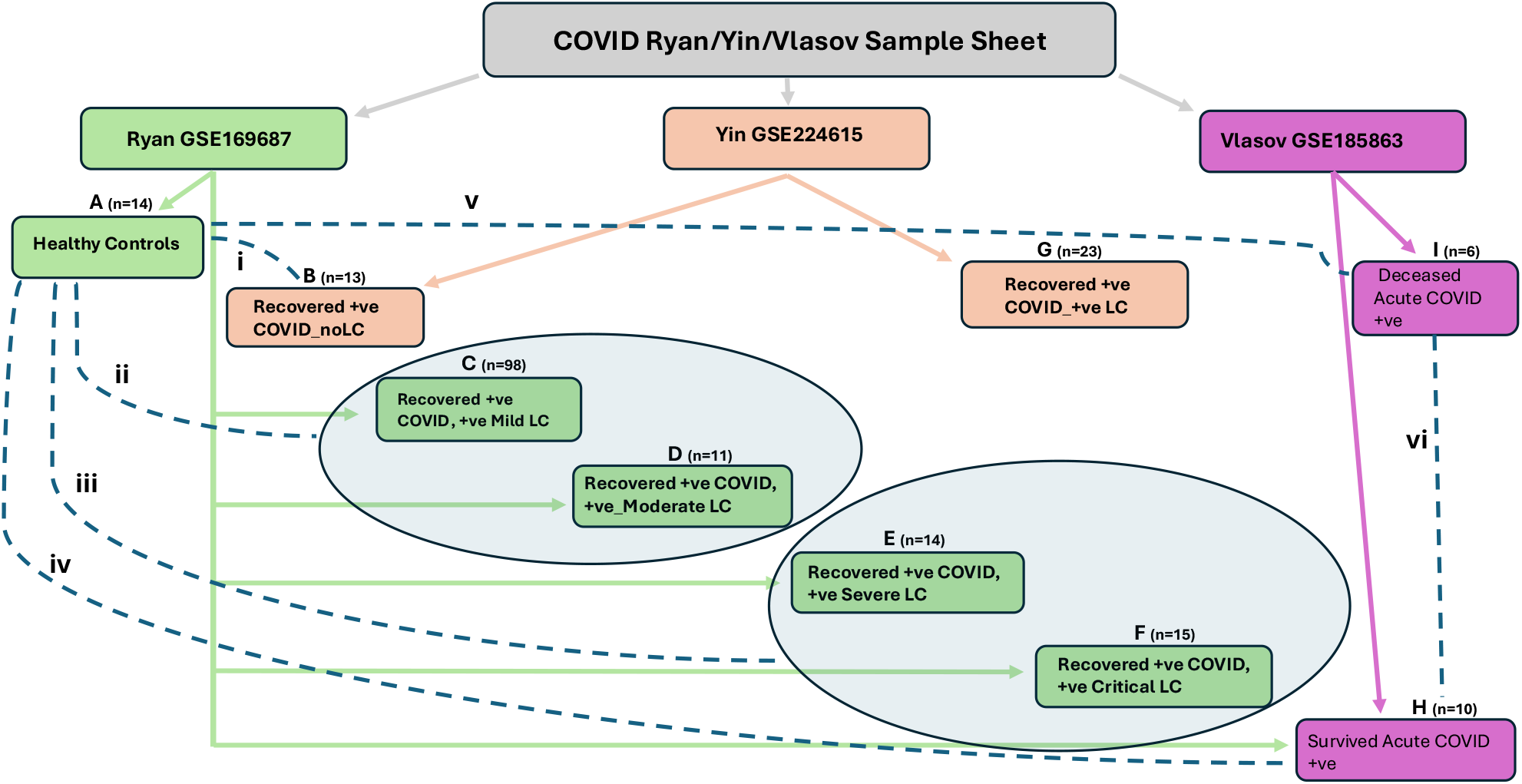
An illustration of IUP’s researcher’s experimental design for inferring DEGs covering a large portion of COVID-19 and post-acute LC pathophysiology for elucidation of intracellular determinants. Gray ovoids were merged during analysis producing C & D as well as E & F categories. Roman numeral on the dashed lines represent the comparisons used for DEGs.

#### Ryan et al. Study Characteristics

This study from Australia enrolled 69 convalescent patients that had provided informed consent between March and April 2020 with SARS-CoV-2 diagnosis confirmed using PCR of nasopharyngeal swabs^14^. Healthy controls (n=14) were defined as individuals with no respiratory symptoms, negative PCR results, no active systemic diseases, and absence of anti-spike or anti-RBD antibodies. This dataset contains negative controls and classified PASC cases grouped as mild, moderate, severe, and critical.

#### Vlasov et al. Study Characteristics

This study which was conducted in Russia included 16 patients of Russian ethnicity aged 40-80 years, all with RT-PCR confirmed SARS-CoV-2 and clinical classification of severe active pneumonia needing ICU-based care^16^. Patients were followed for up to 30 days with blood samples collected at admission. Notably, only male patients were included in the fatal group that underwent an autopsy. This dataset was critical for examining the gene expression profile in acute severe disease as compared to the negative control dataset.

#### Yin et al. Study Characteristics

This prospective observational study enrolled participants in USA with PCR-confirmed SARS-CoV-2 regardless of post-acute symptom presence. Comprehensive assessment of the patients yielded 27 participants categorized as having Long COVID (median age 46 years, 63% female) and 16 as having recovered without sequelae (median age 45.5 years, 44% female) ^15^. The group with Long COVID was not sub-categorized as was the case with the Ryan Et al study. This dataset enabled investigation of molecular differences between recovery patterns and biomarker development for PASC diagnosis.

### Sample Processing and RNA Sequencing

All studies were selected because they contained blood samples in EDTA-anticoagulated tubes with peripheral blood mononuclear cell (PBMC) isolation for RNA sequencing to further reduce confounding factors. The Vlasov et al. study additionally collected PAXgene samples for RNA protection. Although RNA extraction protocols varied across studies with: Ryan et al. using PAXgene blood RNA kit (Qiagen), Vlasov et al. employed TRIzol (Thermo Fisher Scientific), and Yin et al. utilized AllPrep (Qiagen), our uniform processing ensured meaningful integration for downstream analysis. Library preparation similarly varied, with Ryan et al. using Nugen Universal plus total RNA-seq kit, Vlasov et al. employing NEBNext ultra II RNA library prep kit, and Yin et al. utilizing commercial services (Genewiz/Azenta).

### Uniform Bioinformatics Processing Pipeline

We implemented a uniform computational pipeline for processing of all datasets to address confounders that had the potential to obscure biological insights relevant to our main objectives. Figure 2 shows an overview of our analytical pipeline.

**Figure 2:**
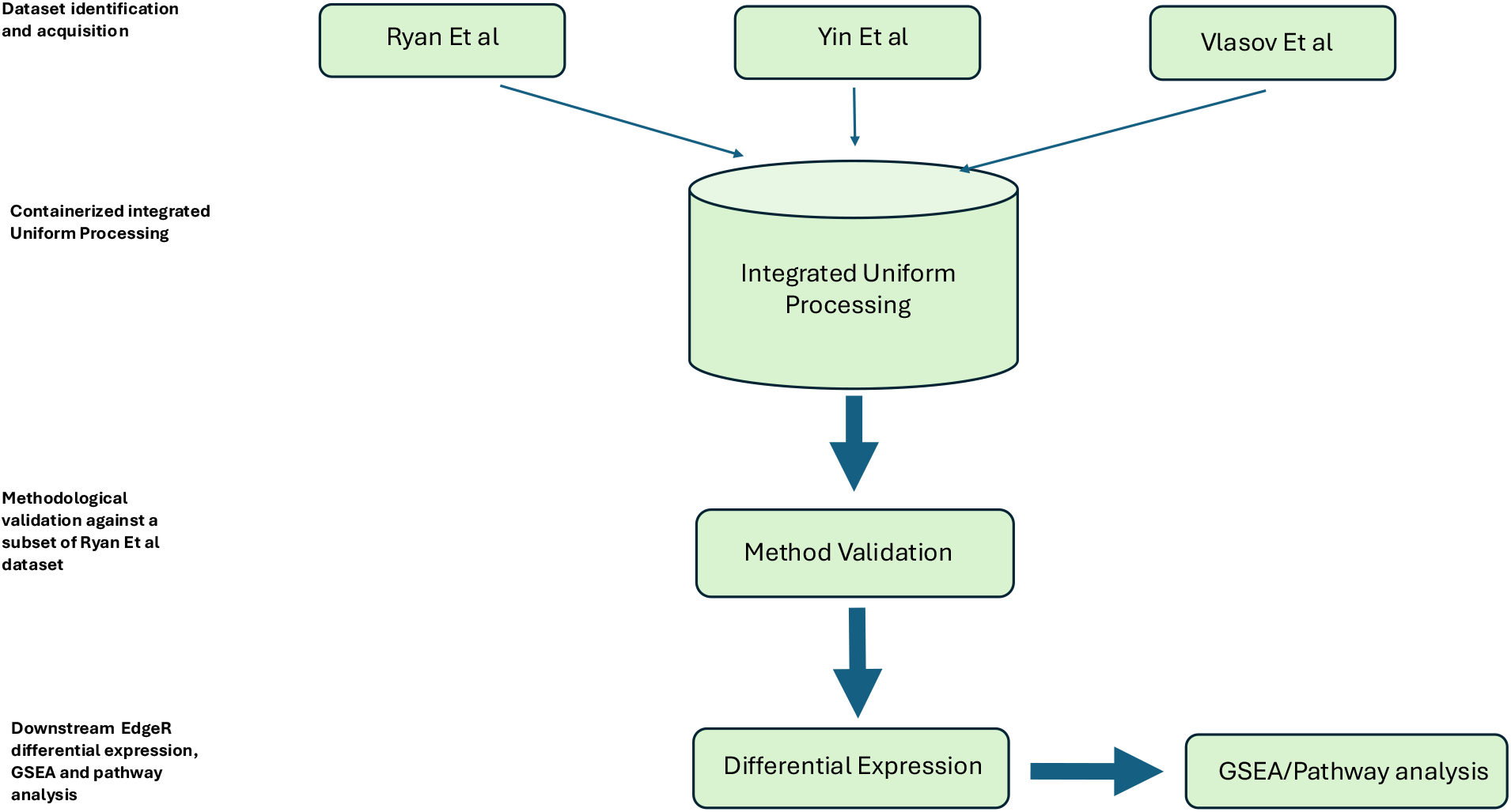
Flow diagram describing the full pipeline of the study methods used in IUP.

We used the GRCh38 primary assembly reference genome (Ensembl release 110) and the GENCODE primary assembly annotation (release 44) as the basis for sequence alignment and quantification. In the beginning stages of our workflow, we derived a transcriptome from the genome and annotation using the GffRead (version 0.12.7)^17^, which extracts transcript sequences from genome annotations. To obtain the sequencing files for samples associated with a study, we downloaded FASTQ files using SRA-toolkit’s fastq-dump utility, specified by the sample’s SRA Run ID (SRR). We used Trim Galore (version 0.6.10)^18^ for quality and adapter trimming on the FASTQ files.

After trimming, we aligned the FASTQ files to the reference genome using STAR (version 2.7.11a)^19^. We provided STAR the “TranscriptomeSAM” setting to translate the alignments into transcript coordinates in an aligned BAM file. We used these BAM files to obtain quantifications using Salmon (version 1.10.1)^20^ subsequently using the gene-level counts from Salmon for downstream analyses.

### Data Processing and Quality Control

Post-alignment processing included sample extraction based on SRR and GSM accession identifiers from the NCBI Sequence Read Archive (SRA) and Gene Expression Omnibus (GEO) to ensure sample identifier synonymity conducive to programmatic sample sheet use. Samples processed in replicates were averaged to generate singular representations and Ensembl gene IDs were mapped to gene symbols using BioMart to ensure unique gene representation thereby eliminating instances of multi-mapping.

Low-count transcripts were filtered out using a 5% expression threshold to drop poorly expressed genes that could introduce noise during downstream analysis. Validation of processing pipeline accuracy was performed by reproducing a subset of Ryan et al. results using identical analytical parameters Figure 3, with Pearson correlation analysis confirming processing fidelity despite differences in alignment algorithms (STAR vs. HiSAT2), Figure 4.

**Figure 3.**
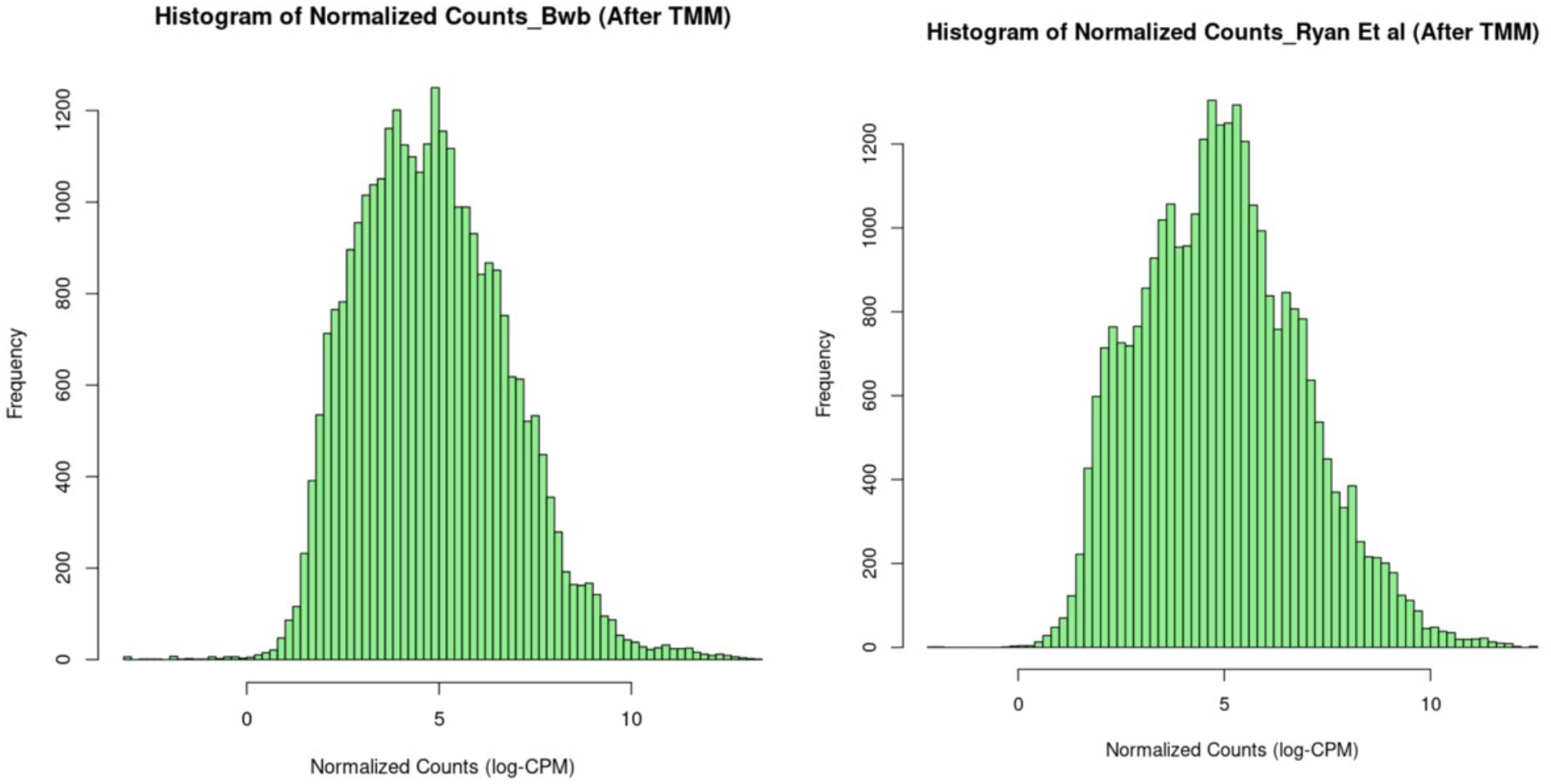
Side-by-side histograms comparing the shape of post-normalization outputs for IUP and Ryan Et al

**Figure 4.**
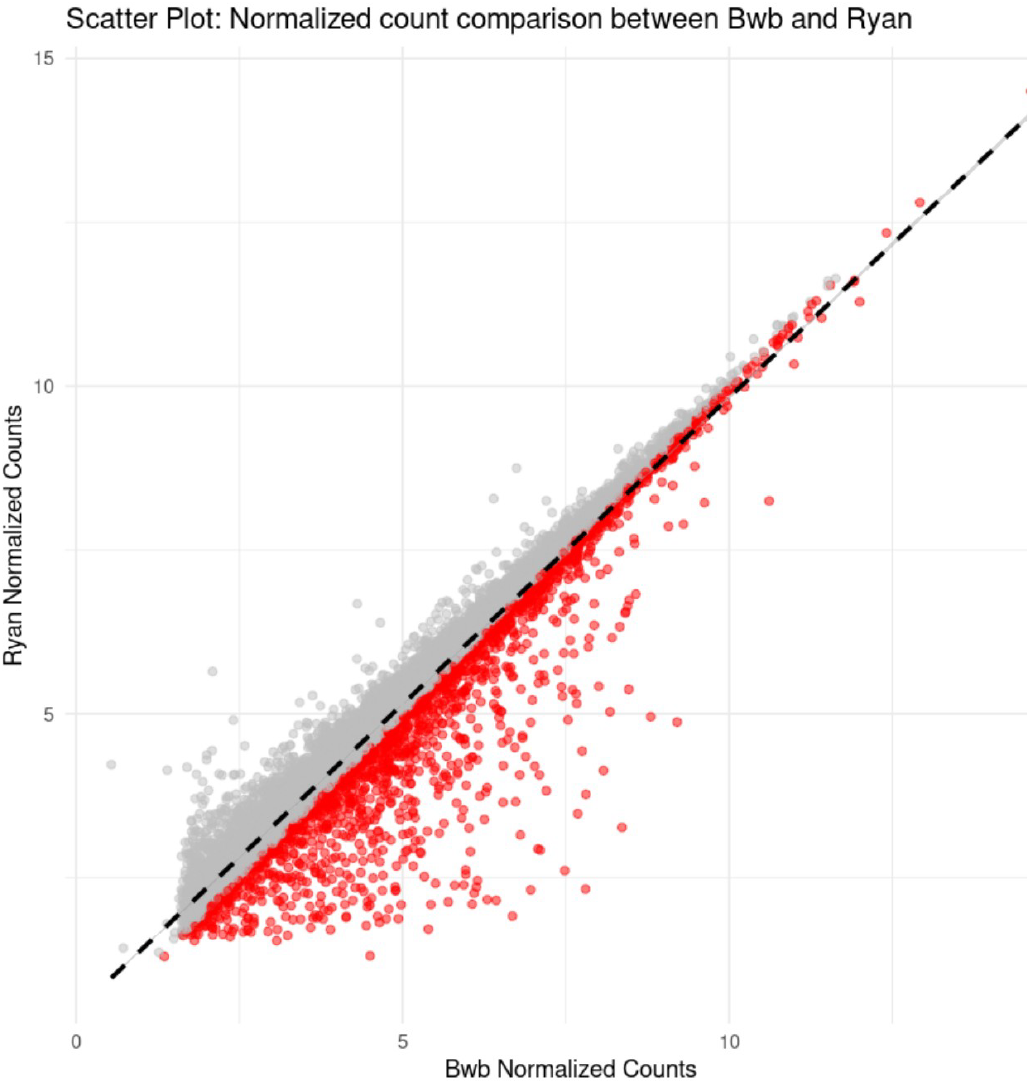
A Scatter plot displaying the overall correlation of the top 1000 genes by expression levels between samples analyzed using Ryan Et al’s (Gray) method versus those produced using IUP’s (Red) method.

**Figure 5.**
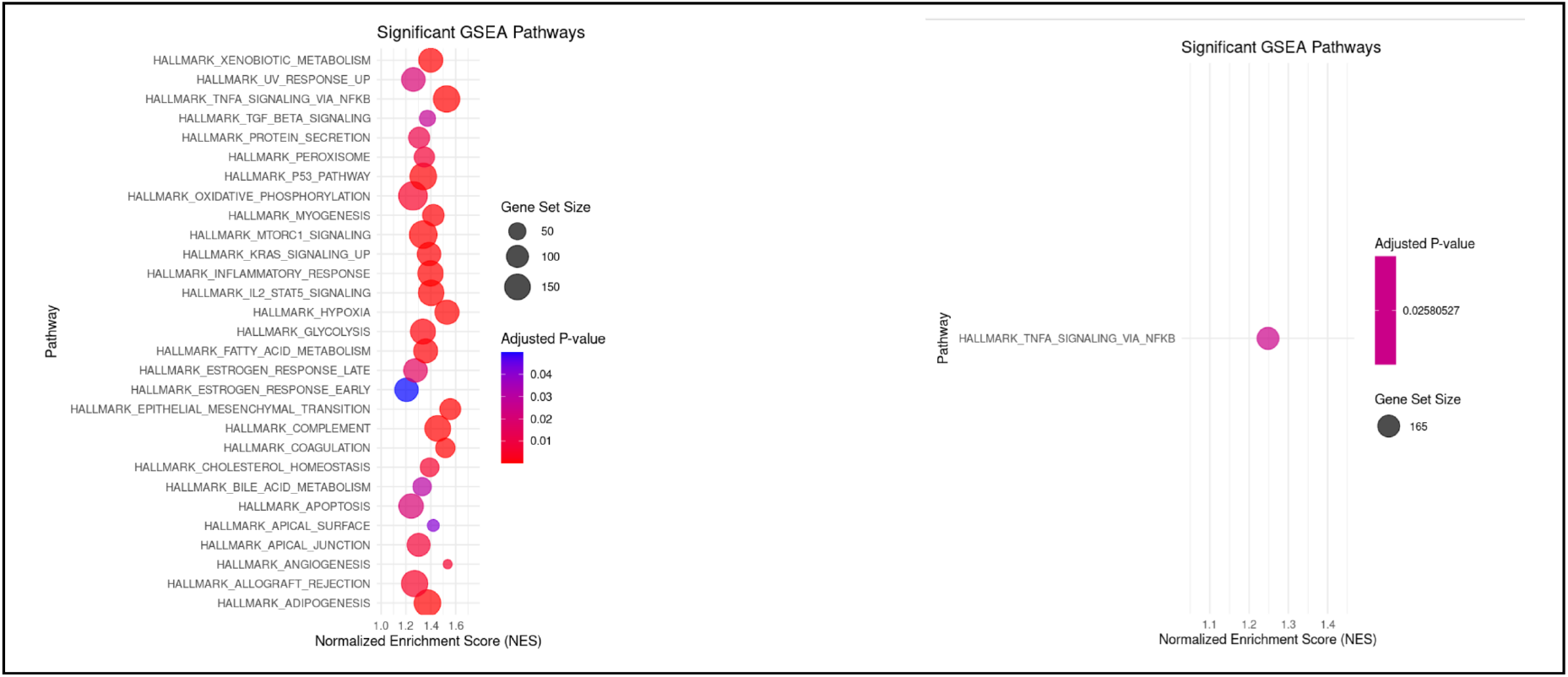
A side-by-side comparison of the enriched pathways using the NES to sort in descending order along with the gene set size displayed by the size of each circle. The adjusted p-values are included with blue and red being the closest to and furthest from 0.05 and red having the smallest value.

### Experimental Design and Comparative Analysis Framework

Our study design had six main comparisons enabling us to systematically investigate pathway-specific molecular signatures across the COVID-19 disease spectrum, specifically targeting identification of gene expression markers and characteristics dominating active severe versus PASC phases of COVID-19. See Figure 1 for a summary of our experimental design:

i. **Group A vs B**: Healthy controls vs. recovered patients with no PASC signs and symptoms (to identify the main post-recovery molecular signatures)
ii. **Group A vs C+D**: Healthy controls vs. mild/moderate PASC (to help us investigate the dominant gene expression pathways as patients enter the PASC phase)
iii. **Group A vs E+F**: Healthy controls vs. severe/critical PASC (to characterize the molecular associations differentially expressed genes with severe/critical PASC)
iv. **Group A vs H**: Healthy controls vs. acute severe COVID-19 survivors (examining enriched gene pathways around resolution of severe active COVID-19)
v. **Group A vs I**: Healthy controls vs. acute severe COVID-19 fatalities (identifying maximal infection related inflammatory pathway activation secondary to COVID-19)
vi. **Group H vs I**: Acute severe survivors vs. fatalities (contrasting the dominant gene expression profiles of pathways related to fatality versus near-fatal cases)

This design enabled systematic investigation of the dominant pathways acute severe disease contrasted with the stratified severity related sequelae experienced in PASC.

### Differential Gene Expression Analysis

Differential expression analysis was performed using edgeR^21^ with similar parameters to Ryan Et al: with false discovery rate (FDR) < 0.05 and log2 fold-change > 1.25. These criteria were selected to ensured identification of robust molecular signatures suitable for pathway analysis and biomarker development due to their stringent thresholds.

### Gene Set Enrichment Analysis (GSEA)

GSEA was performed using MSigDB hallmark gene sets (h.all.v2023.1.Hs.entrez.gmt) downloaded from the Broad Institute and Ensembl gene IDs were converted to Entrez IDs and gene symbols as necessitated. Data preprocessing included removal of NA values and resolution of duplicate FDR values through minimal mathematical perturbation (SD = 1e-8) to maintain dataset integrity and dimensionality while enabling proper ranking for GSEA analysis.

## Results

### Validation of Uniform Processing Pipeline

To ensure analytical validity of our integrated uniform processing, we used our output counts matrices that used STAR aligner to reproduce key findings from a subset of the Ryan et al. study. Using identical analytical parameters but different alignment software (STAR in our case vs. HiSAT2 used by Ryan Et al), we achieved strong correlation in gene expression profiles and identified 15 common differentially expressed genes among the top 50, confirming the robustness of our processing pipeline for downstream pathway analysis (Supplimentary S1).

### Pathway-Specific Molecular Signatures Across COVID-19 Disease Spectrum

Our systematic analysis of six comparative groups revealed distinct molecular signatures and gene-set enrichment that support our three primary findings regarding inflammatory (in active severe COVID-19) and metabolic pathway (PASC) dominance across COVID-19 phases.

### TNF-α/NF-κB Pathway Dominance in Acute Severe COVID-19

Analysis of acute severe COVID-19 cases (Groups H and I) versus healthy controls (Group A) showed predominant enrichment of TNF-α signaling via NF-κB pathways. This inflammatory cascade was most pronounced in cases resulting in fatalities, supporting the premise that hyperinflammatory responses characterize acute severe disease. The pathway enrichment included key inflammatory mediators (IL-6, TNF-α) and anti-apoptotic proteins (BCL2), which is consistent with a cytokine storm pathophysiology and cellular survival responses in acute infection.

**Figure.**
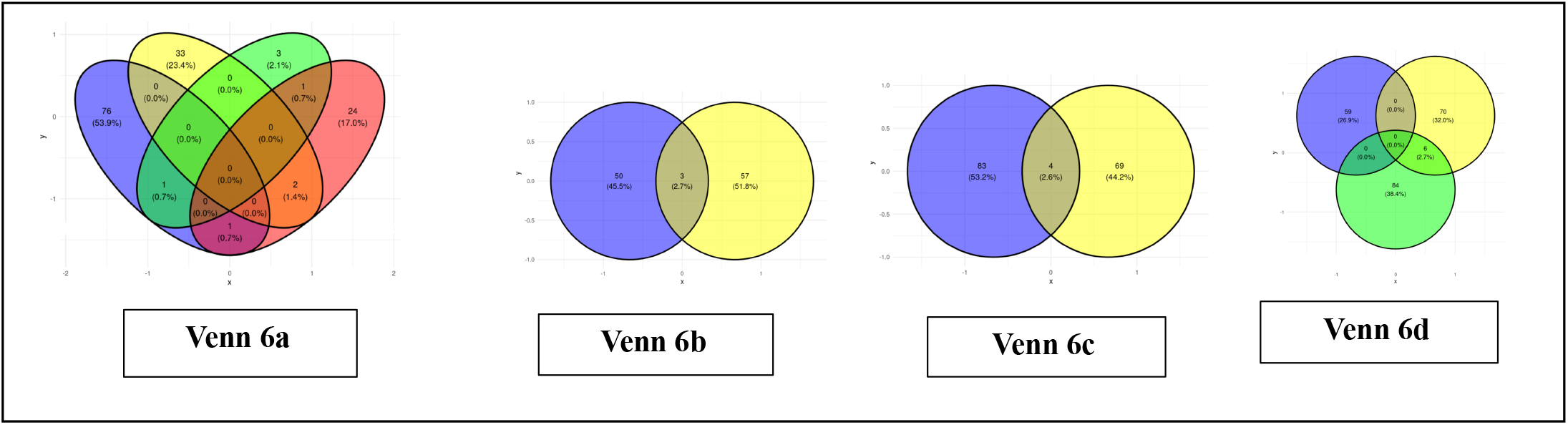
**Venn 6a**: Showcases a Venn diagram with the top four most enriched pathways at the intersection of H versus I based on NES. Note: No intersections were revealed between A versus H or A versus I. **Venn 6b:** This panel is composed of three sets of Venn diagrams showcasing the NES results for each of the comparisons in the Long-Covid datasets. There were two NES enriched pathways that featured in all the comparisons with the A versus B group having an additional enriched pathway (green color).

### Metabolic Pathway Dysregulation in PASC

In contrast, analysis of PASC patients across severity levels (Groups B, C+D, E+F) revealed dominant enrichment of Myc Targets V1 and Oxidative Phosphorylation pathways. This pathway showed sustained activation affecting cell cycle genes (CCND2), glycolytic enzymes (LDHA), and ribosome biogenesis factors (RPL family), suggesting ongoing cellular reprogramming along with metabolic adaptation. Oxidative phosphorylation pathway upregulation may represents compensatory responses to mitochondrial dysfunction, providing a plausible molecular explanation for the characteristic fatigue and post-exertional malaise observed in PASC patients.

**Figure 13.**
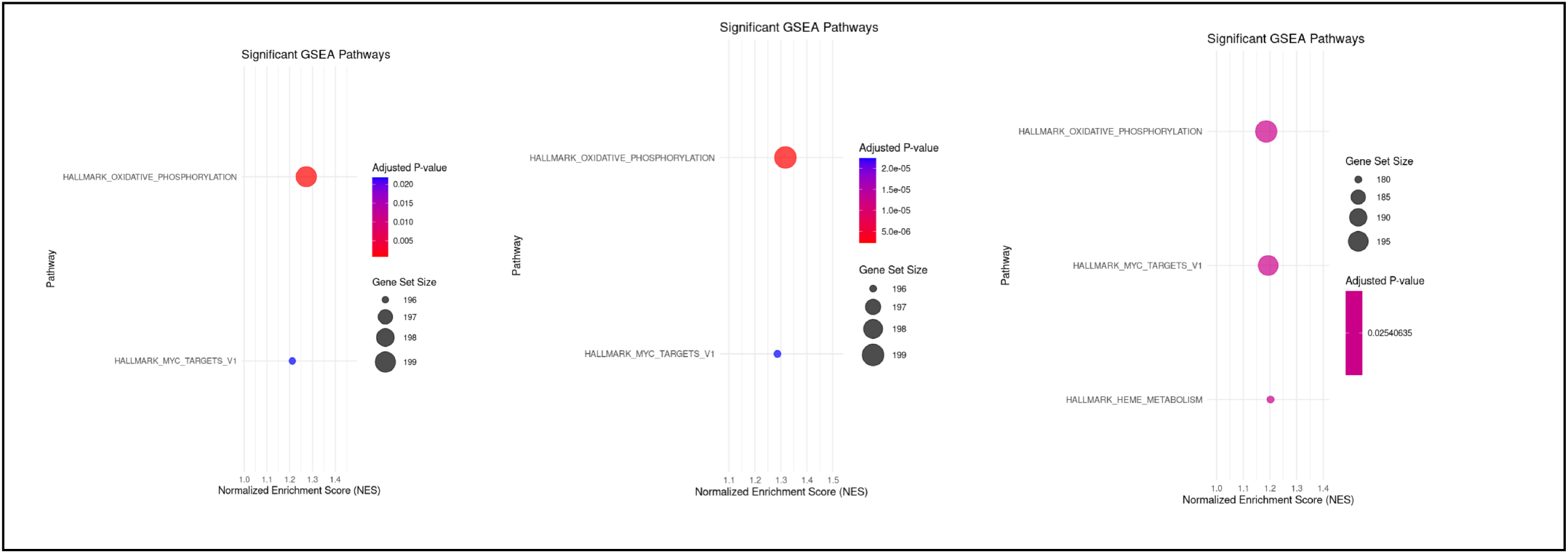
A side-by-side comparison of the enriched pathwaysn using the NES to sort in descending order along with the gene ste size displayed by the size of each circle. The adjusted p-values are included with blue and red being the closest to and furthest from 0.05 and red having the smallest value.

### Progressive Pathway Transition from Acute to Chronic Phases

Our compendium analysis revealed a clear progression from inflammatory to metabolic pathway dominance, from a gene expression vantage point, across disease phases . Even recently recovered patients without PASC symptoms (Group B) showed TNF-α pathway activation, suggesting inflammatory resolution is gradual. However, patients with established PASC demonstrated complete metabolic pathway dominance, indicating a fundamental shift in cellular pathophysiology from acute survival-focused inflammatory responses to chronic adaptation-focused metabolic reprogramming.

### Molecular Biomarkers for Disease Phase Classification

The distinct pathways identified in our analysis provide objective molecular criteria for COVID-19 phase classification, applicable to our third objective. TNF-α/NF-κB pathway gene expression signatures reliably distinguished acute severe disease from other phases, while Myc and oxidative phosphorylation pathway signatures specifically identified PASC patients across severity levels. These find utility in delineating active COVID-19 from the different severities of PASC and in the case of PASC, may offer attractive targets for direct diagnosis which is currently not an option.

These findings establish a molecular framework for precision diagnosis, moving beyond pure subjective symptom-based assessments such as questionnaires and elimination processes, to objective transcriptomic biomarkers. The clear dichotomy between inflammatory and metabolic pathway dominance provides a foundation for developing clinical assays that could enable real-time disease phase monitoring thereby guiding phase-appropriate therapeutic interventions.

Our results demonstrate that integrative analysis of uniformly processed transcriptomic data, from different sources and analytical protocols, can reveal clinically actionable molecular insights, establishing a methodological framework that can be applied to other complex disease processes.

## Discussion and Conclusion

Our analysis based on this compendium of bulk RNA-seq datasets reveals striking dichotomies distinguishing acute severe/critical COVID-19 from post-acute sequelae of COVID-19 (PASC). These findings illuminate fundamentally different mechanisms underlying each phase of the disease, with profound implications for diagnosis, monitoring, and treatment strategies.

### Acute Severe/Critical COVID-19: Inflammatory Dominance

Patients with active severe/critical COVID-19 demonstrate predominant enrichment of the TNF-α signaling via NF-κB pathway, pointing to a cytokine storm survival phenomenon. This hyperinflammatory cascade commences when viral infection triggers TNF-α production through pattern recognition receptors, initiating a devastating positive feedback loop culminating in NF-κB activation, which in turn drives the expression of IL-6, IL-1β, and additional TNF-α, perpetuating the inflammatory spiral that characterizes severe COVID-19 pathophysiology.

This molecular signature correlates well with the known clinical presentation whose hallmarks include rapid onset of dyspnea, fever, hypoxemia, and multi-organ dysfunction, often requiring intensive supportive care including mechanical ventilation, corticosteroids, and immunomodulatory interventions. The dominance of this inflammatory pathway during acute severe disease provides a mechanistic explanation for the clinical efficacy of anti-inflammatory treatments such as dexamethasone and tocilizumab in critically ill patients.

### Post-Acute Sequelae of COVID-19: Metabolic Dysregulation and Repair Dominance

In stark contrast, PASC patients exhibit dominant enrichment in Myc Targets V1 and Oxidative Phosphorylation pathways, indicating a fundamental shift from survival hyperinflammation to an adaptive pathophysiology. This transition suggests that cellular growth, regeneration, and cellular energy metabolism become central drivers in this chronic phase.

The prominence of oxidative phosphorylation pathway enrichment likely reflects mitochondrial dysfunction whose etiology requires further mechanistic elucidation. The upregulation of OXPHOS genes may represent a compensatory attempt to restore mitochondrial homeostasis, potentially explaining the debilitating fatigue characteristic of PASC. This finding aligns with emerging evidence of mitochondrial dysfunction in other post-viral syndromes and chronic fatigue conditions.

The clinical phenotype aligns with this metabolic dysregulation: chronic fatigue with post-exertional malaise, cognitive impairment, and multi-system symptoms necessitating rehabilitation-focused interventions rather than acute inflammatory suppression. The Myc pathway activation suggests ongoing cellular reprogramming and proliferative responses that may contribute to tissue remodeling and potentially maladaptive repair processes.

### Clinical Translation and Therapeutic Implications

The distinct molecular signatures offer objective biomarkers for COVID-19 phase classification, with TNF-α/NF-κB pathway signatures identifying patients at risk of severe disease progression, while OXPHOS and Myc pathway-associated signatures may facilitate PASC diagnosis and severity stratification. This addresses the current reliance on subjective questionnaire-based and elimination-based diagnostic approaches that often delay appropriate care.

Longitudinal monitoring of pathway signature evolution could predict recovery trajectories, thereby identifying patients likely to progress from acute infection to chronic PASC versus those likely to fully recover. Such prognostic capabilities would enable early intervention strategies and inform patient counseling regarding expected disease courses.

These findings support fundamentally different treatment paradigms for each phase. Acute severe COVID-19 benefits from anti-inflammatory interventions targeting the dominant TNF-α/NF-κB pathway, consistent with the proven eSicacy of corticosteroids and immunomodulators. Conversely, PASC management should prioritize mitochondrial function restoration and cellular repair mechanisms. Therapeutic targets may include mitochondrial support compounds, metabolic modulators, and interventions targeting cellular energetics, representing a paradigm shift from inflammatory suppression to metabolic restoration and adaptation.

### Future Directions

Our findings exemplify how compendium-based transcriptomic analysis can reveal disease-specific molecular mechanisms with immediate clinical relevance. The clear progression from inflammatory to metabolic pathway dominance provides a framework for precision, phase-appropriate interventions-a critical step towards personalized medicine in COVID-19 care.

Longitudinal pathway monitoring studies could also help define biomarker panels for clinical implementation, while phase-specific therapeutic trials could transform COVID-19 and PASC management to include the molecular mechanisms of disease in patient care. The flexible analytical framework demonstrated in this work offers broad adaptable applicability across many disease processes potentially accelerating personalized medicine development through AI-enabled analysis.

## Data Availability

All datasets can be found in the NCBI repository (https://www.ncbi.nlm.nih.gov/geo/) via their respective accession numbers as follows: Ryan Et al – GSE169687, Yin Et al – GSE224615, and Vlasov Et al – GSE185863.

